# Separate to operate: the centriole-free inner core of the centrosome regulates the assembly of the intranuclear spindle in *Toxoplasma gondii*

**DOI:** 10.1101/2022.01.05.475174

**Authors:** Ramiro Tomasina, Érica S. Martins-Duarte, Philippe Bastin, Mathieu Gissot, María E. Francia

## Abstract

Centrosomes are the main microtubule-organizing center of the cell. They are normally formed by two centrioles, embedded in a cloud of proteins known as pericentriolar material (PCM). The PCM ascribes centrioles with their microtubule nucleation capacity. *Toxoplasma gondii*, the causative agent of toxoplasmosis, divides by endodyogeny. Successful cell division is critical for pathogenesis. The centrosome, one of the microtubule organizing centers of the cell, plays central roles in orchestrating the temporal and physical coordination of major organelle segregation and daughter cell formation. The *Toxoplasma* centrosome is constituted by two domains; an outer core, distal from the nucleus, and an inner core, proximal to the nucleus. This dual organization has been proposed to underlie *T. gondii*’s cell division plasticity. Homeostasis of the outer core has been shown to be critical for the proper assembly of the daughter cells. However, the role of the inner core remains undeciphered. Here, we focus on understanding the function of the inner core by studying the dynamics and role of its only known molecular marker; TgCEP250L1. We show that upon conditional degradation of TgCEP250L1, parasites are unable to survive. Mutants exhibit nuclear segregation defects, whilst normally forming daughter cells. In addition, the rest of the centrosome, defined by the position of the centrioles, disconnects from the nucleus. We explore the structural defects underlying these phenotypes by high resolution microscopy. We show that TgCEP250L1’s location is dynamic and encompasses the formation of the mitotic spindle. Moreover, we show that in the absence of TgCEP250L1, the microtubule binding protein TgEB1, fails to translocate from the nucleus to the mitotic spindle, while polyploid nuclei accumulate. Overall, our data supports a model in which the inner core of the *T. gondii* centrosome critically participates in cell division by directly impacting the formation or stability of the mitotic spindle.

## INTRODUCTION

Centrosomes are the main microtubule-organizing centers (MTOC) of the cell. In mammalian cells, the centrosome is formed by two microtubule-based barrels, known as centrioles, which display a highly conserved, nine fold radial symmetry of triplet microtubules. Centrioles reside within a complex matrix of proteins, collectively known as the pericentriolar material (PCM). The PCM ascribes centrioles with their microtubule nucleation capacity. The centrosome’s microtubule organization capacity plays pivotal roles in cellular life, impacting cell shape and polarity, organizing the formation of motile structures, and participating in karyokinesis.

The phylum Apicomplexa is a large group of protozoan parasites, consisting of more than 6000 species (1). Apicomplexans cause important human and animal diseases including toxoplasmosis, malaria, neosporosis and cryptosporidiosis. Toxoplasmosis is caused by *Toxoplasma gondii*, arguably the most successful parasitic organism of warm-blooded animals in the world. It is estimated that around 30% of the human population is infected with this parasite. The most severe outcomes by *T. gondii* infection are due to reactivation of chronic infection, primo infections in immunocompromised individuals, and congenital transmission (2, 3). The latter could lead to miscarriage or irreversible sequelae for the newborn.

*T. gondii* actively invades virtually any nucleated cell. Once inside the cell, it replicates, augmenting its numbers rapidly and exponentially, eventually causing host cell lysis. Newly released parasites can subsequently invade healthy neighboring cells perpetuating the infection. The fast dividing form of the parasites, known as the tachyzoite, follows a cell division scheme known as endodyogeny. Endodyogeny consists of a semi-closed nuclear mitosis - i.e. no appreciable chromatin condensation nor nuclear envelope breakdown occur - concomitant with the assembly of two daughter cells within the mother cell (4).

Tachyzoites bear two MTOCs: the Apical Polar Ring (APR) and the centrosome. The APR is involved in nucleating the cortical microtubules that shape and permit parasite motility(5). The centrosome in *T. gondii* has been shown to orchestrate the temporal and spatial coordination of nuclear mitosis and daughter cell formation (6, 7). On one hand, it nucleates the mitotic spindle microtubules, impacting chromatin organization and nuclear segregation. On the other hand, the centrosome organizes the seeds of new cells by physically positioning the offspring’s APR, thereby spatially and temporally linking new daughter cell formation with nuclear content segregation(8).

Recently, the centrosome in *T. gondii* was shown to be constituted by three distinct protein localization domains. Initially, an outer core, distal from the nucleus, and an inner core, proximal to the nucleus, were identified(9). These domains were described based on the localization of centrosomal protein homologs. A homolog of Centrin1, an EF-hand calcium binding protein and a *bona fide* marker of centrioles in many species, was shown to localize at the outer core. Likewise, SAS6, a protein involved in forming the cartwheel, the structure responsible for ascribing centrioles with their characteristic geometry, co-localizes with Centrin1. Sfi1 and γ-tubulin homologs are also found at the outer core. Centrioles likely reside within the outer domain. However, this has not been experimentally validated. On the other hand, TgCEP250L1 (TgME49_290620), a distant homolog of CEP250, a centrosomal protein involved in centriole cohesion, localizes to the inner core exclusively (9, 10). An additional CEP250 homolog localizes to both the inner and outer cores (9). Experimental manipulation of TgCEP250 causes physical separation of the cores and the concomitant dysregulation of cytosolic and nuclear events during cell division. This protein has been proposed to bridge cohesion between the two opposite cores (10). More recently, a third protein localization domain, in between the initially described outer and inner cores, was identified. The “middle” core houses TgCEP530; a mutant of this protein loses synchrony between cytokinesis and karyokinesis and exhibits outer core fragmentation, indicating that this domain also plays a role in the regulation of cell division and is important for centrosomal homeostasis (11).

While the outer core has been proposed to regulate aspects of daughter cell formation, the inner core is intuitively associated with nuclear events. However, how the different domains of the centrosome interplay to coordinate aspects of cell division remains poorly understood. In particular, the role played by the inner core remains experimentally unexplored. To gain further insight into these aspects of *T. gondii*’s cell division, here we focus on the characterization of TgCep250L1’s function, as a proxy to the role of the centrosomal inner core in this parasite

## MATERIALS AND METHODS

### Parasite culture

*T. gondii* tachyzoites of the RHΔKu80 strain expressing the Tir1 receptor,(12) were maintained in VERO cells, grown in Dulbecco’s modified Eagle’s medium (DMEM; Gibco, St. Louis, USA) supplemented with 10% fetal bovine serum (FBS; Gibco, St. Louis, USA), 4 mM of L-Glutamine (Gibco, St. Louis, USA), 200 U/ml of penicillin and 200 μg/ml of streptomycin (Gibco, St. Louis, USA). Cultures were kept at 37°C and 5% CO2.

### Generation and preliminary characterization of the TgCep250L1 inducible knockdown strain

TgCEP250L1-mAID-3HA was generated in the RHΔku80 Tir1 strain background (13). Briefly, a PCR product of the mAID-3HA sequence bearing 35bp of homology to either end of the TgCEP250L1 gene stop codon, was amplified using the primers 3 and 4 (Supplementary Table S1). A plasmid coding for an RNA guide targeting the 3’ end of the TgCEP250L1 gene and SpCas9 was generated by mutagenesis using primers 1 and 2 (Supplementary Table S1), then 50 μg of pSagCas9 vector and 10μg of PCR product were transfected into 5.0 × 10^7^ parasites using a BTX 600 electroporator, following previously published protocols (14). The pSagCas9 vector (15) was kindly provided by Dr. David Sibley.

Successful insertion of the 3HA-mAID sequence was monitored by PCR using primers 5 to 7 (Supplementary Table S1) according to the schematic shown in Supplementary Figure 1A. Clonal cell lines were obtained by limiting dilution. Protein degradation was triggered by addition of 0.5 mM indole-acetic acid (IAA; Sigma-Aldrich) to the growth medium.

### Plaque and Intracellular Growth Assays

For plaque assay, two hundred parasites of the TgCEP250L1-mAID-3HA strain were inoculated on Human Foreskin Fibroblasts cells (HFF; kindly provided by Dr. Sebastian Lourido) previously grown to confluency on six well plates, and kept for seven days in presence or absence of 0.5 mM IAA. Wells were then fixed with methanol and stained with Crystal violet for plaque visualization. Intracellular growth was determined by immunofluorescence assay (IFA; protocol specified below) labeling parasites with the pellicle marker anti-IMC1 and DAPI (nuclear and apicoplast DNA marker). Assays were done by infecting confluent HFF cells grown on 13mm coverslips, with 1,000 parasites. Parasites were allowed to invade and grow for 2 hours prior to IAA addition to the media when appropriate. Note that the medium was changed at the same time for parasites grown under control conditions (no IAA). Parasites were allowed to grow for an additional 24 hours, prior to fixation and processing. Assays were done in triplicate. Quantification was done by determining the number of parasites per vacuole in 35 randomly acquired fields at an Olympus epifluorescence microscope.

### Western Blotting (WB)

Total proteins were extracted from 1.0 × 10^8^ parasites grown on media supplemented with 0.5 mM IAA for different times, or as indicated in the figure legends. Total proteins were extracted by resuspending the cell pellet in Laemmli buffer and boiling for 5 minutes. Protein samples were ran on a 10% SDS-polyacrylamide gel for 2 hours, and transferred onto a nitrocellulose membrane overnight. Primary and secondary antibody incubations were done in 5% milk/PBS using rabbit anti-HA at 1:500 (Cell Signaling, Cat #: 3724S) and anti-Rabbit HRP at 1:10,000 (BioRad, Cat #: 1721017). Images were obtained using an ImageQuant 800 Western blot imaging system (Amersham) exposing for a total of 30 seconds.

### Optical microscopy

Immunofluorescence assay (IFA) was performed as previously reported (14). In short, HFF cells were grown in coverslips and inoculated with parasites. Depending on the assay, intracellular parasites were fixed at different times using methanol for 5 minutes at −20°C. Primary antibodies used were: mouse anti-centrin at 1:1,000 (Cell Signaling, Cat #: 04-1624), rabbit anti-HA at 1:200 (Cell Signaling, Cat #: 3724S), mouse anti-IMC-1(16), at 1:500 (Kindly provided by Dr. Gary Ward, University of Vermont), guinea pig anti-TgEB1 (17) at 1:3,000 (kindly provided by Dr. Marc-Jan Gubbels, Boston College), rabbit anti TgH2Bz (18, 19) at 1:3,000 (kindly provided by Dr. Sergio Angel, INTECH-Chascomus), guinea pig anti-NDC80(20) at 1: 2,000 (kindly provided by Dr. Marc-Jan Gubbels, Boston College), mouse anti-γ-tubulin at 1:1,000 (Cell Signaling, Cat #: 5886), and mouse anti-Acetylated tubulin at 1:1,000 (Sigma, Cat #: T7451). Goat anti-rabbit Alexa Fluor 405 (Invitrogen, Cat. # A-31556), Goat anti-rat Alexa Fluor 488 (Invitrogen, Cat. # A-11006), Goat anti-mouse Alexa Fluor 488. (Invitrogen, Cat. # A28175), Goat anti-rabbit Alexa Fluor 488 (Invitrogen, Cat. # A-11008), Goat anti-guinea pig Alexa Fluor 594 (Invitrogen, Cat. #A-11076), Goat anti-rabbit Alexa Fluor 647 (Invitrogen, Cat. # A27040) and Goat anti-mouse Alexa Fluor 647 (Invitrogen, Cat. # A-21235) were all used at dilution of 1:2,000. Coverslips were mounted onto glycerol or fluoroshield with DAPI.

Ultrastructure Expansion Microscopy (UExp) was performed following Dos Santos Pacheco & Soldati-Favre without modifications (21). Primary and secondary antibodies were used twice as concentrated for UExM experiments than specified for IFA above.

All images were acquired using a Zeiss confocal LSM880 microscope using a Plan-Apochromat 63×/1.40 oil objective. All images were acquired and processed using the Zeiss ZEN blue edition v. 2.0 software. All images were deconvolved using Huygens Professional version 19.10.0p2 64b (Scientific Volume Imaging, The Netherlands, http://svi.nl).

### Transmission Electron microscopy (TEM)

TEM sample preparation was done following previously published protocols (22). In short, intracellular parasites were fixed in 2.5 % glutaraldehyde/0.1M sodium phosphate buffer for 2 h at room temperature. The fixative solution was washed out 3 times with 0.1M sodium phosphate buffer and infected cells were post-fixed with 1% OsO_4_. Dehydration was done sequentially incubating samples in 30%, 50%, 70%, 90% and pure acetone for 10 minutes each. Embedding was done using epoxy resin (PolyBed resin, Polyscience Inc., Warrington, PA, USA). Ultrathin sections were obtained, stained, and observed in a transmission electron microscope.

### Flow Cytometry

10^6^ tachyzoites were fixed in 70% (v/v) ethanol overnight at −20°C, washed and resuspended in phosphate buffered saline (PBS) and stained using propidium iodide (PI) at a final concentration of 0.2 mg/ml, and 250U of RNase A (ThermoFisher, #EN0531) for 30 min at room temperature. DNA content was measured based on PI fluorescence using a 488 nm argon laser on a BD Accuri™ C6 Plus Flow Cytometer (Becton-Dickinson, San Jose, CA). Fluorescence was collected in linear mode for 10,000 events per condition.

## Results

To assess the role of the centrosomal inner core of *T. gondii*, we generated a knock down strain of its only identified marker: TgCep250L1 (**Fig. 1A**). We generated TgCEP250L1-mAID-3HA strain by inserting a mini-auxin inducible degron sequence (mAID) (12), followed by a triple hemagglutinin epitope tag (3HA), in frame with the TgCEP250L1’s coding sequence (**Supplementary Fig. 1A**). Successful modification of the locus was corroborated by PCR (**Supplementary Fig. 1B**). Immunofluorescence assays (IFAs) using anti-HA antibodies to visualize TgCEP250L1, and anti-Centrin as a proxy for the centrosome position, show that the TgCEP250L1-mAID-HA fusion correctly localizes to the organelle (**Fig. 1B**).

**Figure 1.**
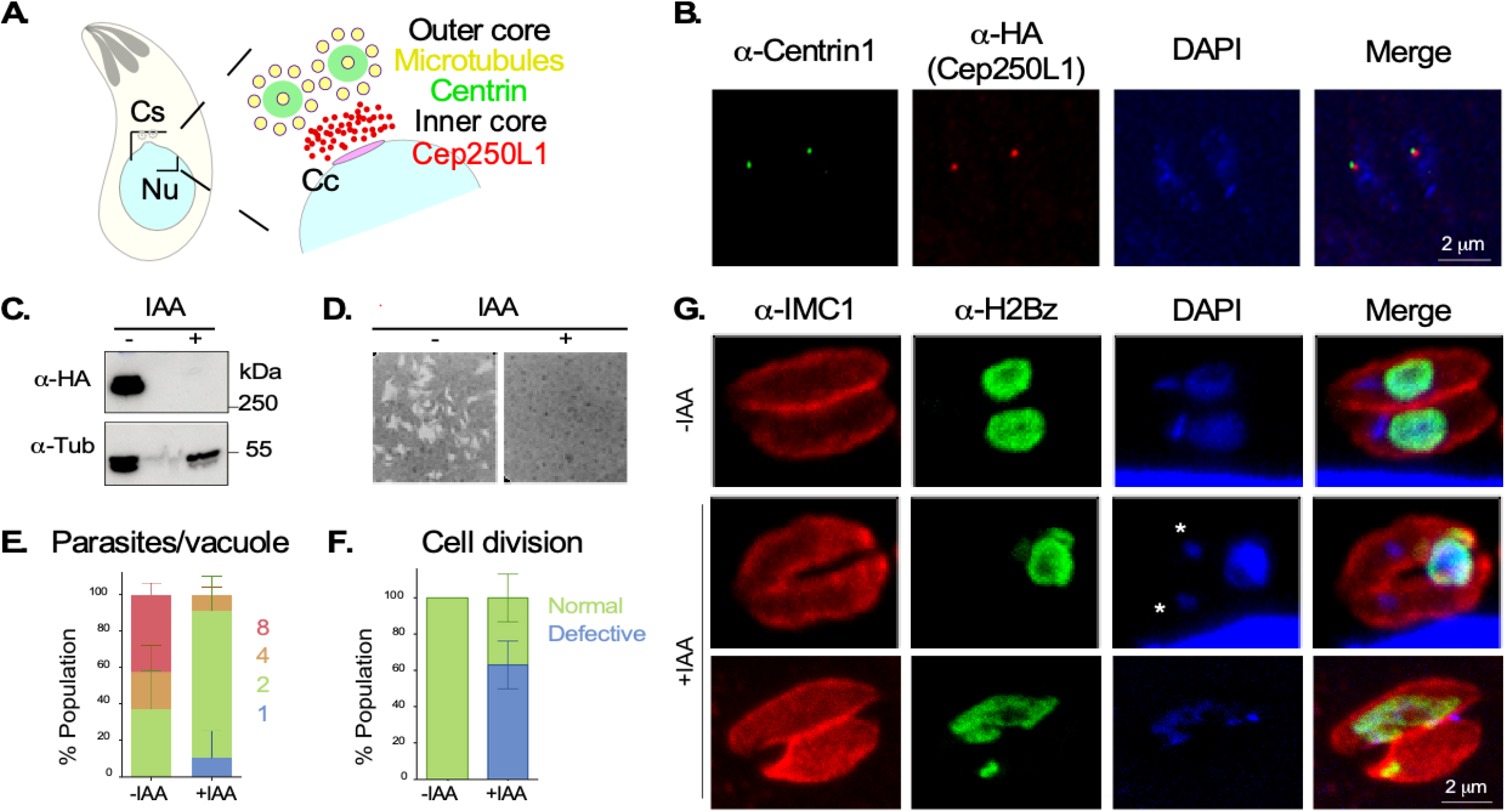
Conditional Knockdown of TgCEP250L1 causes nuclear segregation defects. (A) Schematic representation of the bipartite centrosome of *Toxoplasma gondii*. The centrosome (Cs) of *T. gondii is* organized in two core domains. The outer core houses proteins such as Centrin1, and has been proposed to house two parallel centrioles displaying a nine-fold symmetry of single microtubules. The relative position of TgCEP250L1 is shown. The centrocone (Cc) is a nuclear (Nu) envelope elaboration apposed to the centrosome. (**B) The fusion protein TgCEP250L1-mAID-3HA correctly localizes to the centrosome.** An immunofluorescence assay (IFA) of TgCEP250L1-mAID-HA parasites stained with anti-centrin1 (green), anti-HA antibody for TgCEP250L1-mAID-3HA (red) and DAPI (blue) is shown. (**C) TgCEP250L1-mAID-3HA rapidly degrades upon addition of IAA to the growth media.** Western blot analysis of total protein extracts from the TgCEP250L1-mAID-HA strain treated or not with IAA for 24hs. (**D) TgCEP250L1 is required for tachyzoite proliferation.** Human foreskin fibroblasts monolayers were infected with equal number of TgCEP250L1-mAID-HA and treated, or not, as indicated, for a week. Note that while untreated parasites (-IAA) are able to generate lysis plaques in the monolayer, treated parasites (+IAA) do not proliferate. **(E) TgCEP250L1 knockdown stalls parasite replication.** The number of TgCEP250L1-mAID-HA parasites per vacuole, treated as indicated, was quantified. Data shown are the average of three independent experiments. Error bars represent the standard deviation. (**F) TgCEP250L1 knockdown yields parasites displaying aberrant cell division phenotypes**. The number of parasites displaying miss-segregated or no nuclei were quantified by IFA. Data plotted are the average of three independent experiments. Error bars represent the standard deviation. (**G) TgCEP250L1 depletion causes nuclear missegregation.** TgCEP250L1-mAID-HA parasites, treated as indicated for 24hs, were stained with anti-IMC1 (red; pellicle marker), anti-H2Bz (green; nucleus), and DAPI (blue; DNA marker). Asterisk mark the apicoplast DNA.

Degradation of TgCEP250L1-mAID-HA is rapidly triggered by the exogenous addition of an auxin analogue (indole acetic acid; IAA) to the growth media. Protein knockdown was observed by WB as early as 30 minutes following IAA addition (**Supplementary Fig. 1C**), becoming undetectable in 2 hours **(Fig. 1C and Supplementary Fig. 1C**). Likewise, TgCEP250L1-mAID-3HA readily became undetectable by IFA (**Supplementary Fig. 1C**).

We asked whether TgCEP250L1 plays a role in parasite survival. For this, we pursued plaque assays, whereby we assayed the ability of parasites to survive long-term upon initial protein knockdown. Protein knockdown was triggered at the beginning of the assay, and the ability of parasites to lyse a host cell monolayer was assayed following a weeklong incubation. These assays clearly show that parasites are unable to form plaques, suggesting that TgCEP250L1 is essential for division (**Fig. 1D**).

To understand the mechanisms of parasite death we analyzed parasite growth *in vitro*. For this, we quantified the number of parasites per vacuole upon 24 hours of protein knockdown. The vast majority (80%) of mutant parasites presented two parasites per vacuole, while control parasites (no IAA addition) exhibited 4 to 8 parasites per vacuole (~55%). Only 40% of the control exhibited 2 parasites per vacuole. No vacuoles of 8 parasites were observed in the IAA treated condition (**Fig. 1E**).

In order to understand the underlying defects giving rise to the mutant’s growth arrest, we performed IFA assays after TgCEP250L1 knockdown using anti-IMC1 to label the inner membrane complex, a structure that scaffolds the emerging daughter cells (23); anti-TgH2Bz, to label a histone variant (as a nuclear marker) (18) and DAPI (a DNA labeling dye). While TgH2Bz is specific to the nucleus, DAPI labels not only nuclear chromatin but also the apicoplast genome. This secondary endosymbiont, present in most apicomplexans, bears its own DNA (24). Twenty-four hours following TgCEP250L1 knockdown, forty percent of parasites appear normal whereby each parasite contains a single nucleus. In contrast, 60% of parasites exhibit nuclear segregation defects (**Fig. 1F and G**). Among these defective parasites, 20% of the vacuoles exhibit an individual containing an enlarged nucleus and individuals with either a minimal fragment or no detectable nuclear content. On the other hand, 40% of the vacuoles contain parasites that are seemingly “empty.” The nucleus does not segregate into either one of the forming cells and remains excluded from the newly formed parasites (**Fig. 1G**).

We analyzed whether the nuclear segregation defects are linked to a defect in DNA replication. For this, we assayed chromatin-associated proteins that vary along with ploidy. We assessed the parasites’ kinetochore marker NDC80 (20) or the centromeric histone CenH3 (25). When observed by IFA, antibodies against either one of these proteins label a punctate structure at the nuclear periphery that exists either as a single entity in non-dividing parasites or as a duplicated dot in dividing and segregating nuclei. Quantification of the number of kinetochore/centromeres per nucleus shows that the untreated population exhibits the expected bimodal distribution of 1 or 2 dots. IAA-treated parasites, not only display 1 and 2 dots, but also accumulate nuclei with no NDC80/CenH3 signal (**Supplementary Fig. 2A and 2B**). Additionally, we analyzed the DNA content of propidium iodide labeled parasites by flow cytometry. Untreated parasites exhibit a typical predominant 1N peak, with a small portion of the population exhibiting an expected 1.8N ploidy (26). Overall, we did not detect a significant accumulation of excessive DNA content (> 1.8N), nor parasites exhibiting an excessive number of kinetochores/centromeres per nucleus (>2) in the IAA treated population. This suggests that DNA replication progresses. However, partitioning of nuclear content into daughter cells is unequal.

To better understand the impact of the lack of TgCEP250L1 in *T. gondii* division, we further investigated the ultrastructure of TgCEP250L1-mAID-3HA upon TgCEP250L1 knockdown, by transmission electron microscopy (**Fig. 2**). During cell division, many of the pre-existent mother cell organelles are duplicated and segregated into daughter cells (**Fig. 2A**). These include the centrosome, the nucleus, the apicoplast, the mitochondria and the endoplasmic reticulum, among others (27, 28) (**Fig. 2B**). In the IAA treated population, we observed vacuoles bearing parasites able to correctly form daughter cell scaffolds (**Fig. 2C and D**). Newly formed cells display correctly segregated organelles such as the mitochondria and the apicoplast (insets, **Fig. 2C and D**). Consistent with our observation by IFA (**Fig, 1G**), in several dividing parasites the nuclei appears to be positioned away from the site of daughter cell assembly (**Fig. 2C and D**).

**Figure 2.**
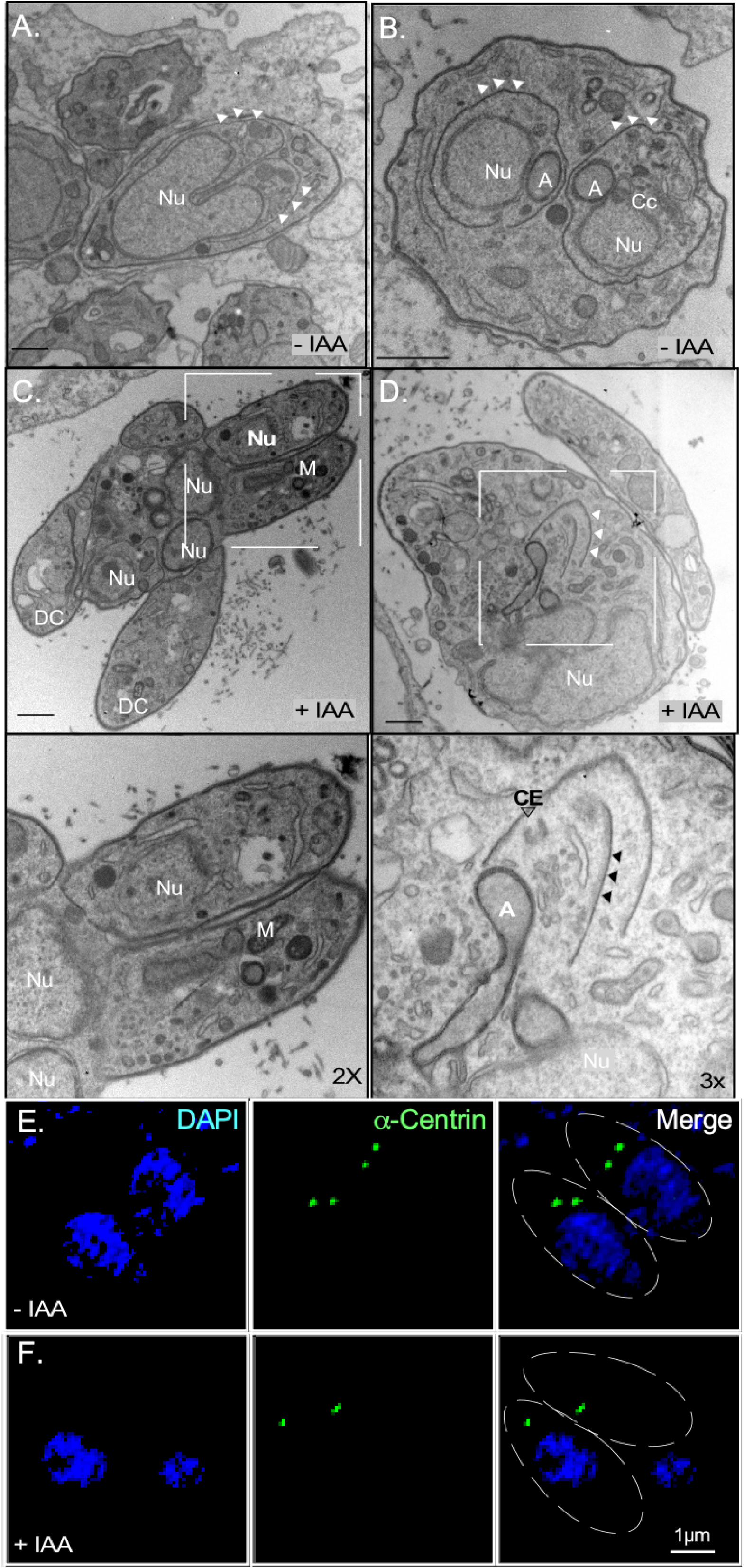
TgCEP250L1 is required for nuclear segregation. **(A**) Electron micrograph **of a *T. gondii* parasite dividing by endodyogeny (longitudinal view**). The image shows the assemblage of two daughter cells scaffolds within a mother cell with the mitotic nucleus (Nu) segregating to each daughter cell. White arrowheads indicate the daughter cells’ inner membrane complex.(**B) Ultrastructure of *T. gondii* parasites upon 24hs of TgCep250L1’s knockdown reveals proper mitochondria segregation.** The *T. gondii* mitochondrion (M) segregates properly. However, several nuclei (Nu) remain excluded from the individual daughter cells. The inset shows a closer view of two of the daughter cells, whereby it can be appreciated one daughter cell inherited a mitochondrion profile (M)**. (C) Electron micrograph of untreated *T. gondii* parasite dividing by endodyogeny (transversal view**). Two daughters’ cells are assembled within a mother cell. A duplicated nucleus (Nu) is packed into each daughter cell. The apicoplast (A) divides by association with the centrosome, and hence segregates physically adjacent to the nucleus (Nu) and the centrocone (Cc). White arrowheads indicate the inner membrane complex of the daughter’s cells. (**D) Ultrastructure of *T. gondii* parasites upon 24hs of TgCEP250L1 knockdown reveals proper apicoplast segregation.** The inner membrane complex of an assembling daughter cell is shown (white arrowheads). The apicoplast (A), a centriole (CE, empty arrowhead, inset) and the striated fiber(8) (black arrowheads, inset) are positioned as expected, and are associated to the daughter cell scaffold. Note that nucleus (Nu), however, remains physically distant from the site of daughter cell assembly. **(E) The centrosome localizes apposed to the nucleus in presence of TgCEP250L1.** IFA using anti-centrin1 marks the position of the outer core of the centrosome (green) in untreated parasites. Approximate parasite position is shown as dotted lines on the merge panel. (**F) Upon TgCep250L1’s knockdown, the centrosome (green) distances from the nucleus (blue**). IFA using anti-centrin1 marks the position of the outer core of the centrosome (green) in IAA treated parasites for 6 hours. Approximate parasite position is shown as dotted lines on the merge panel.

Daughter cell scaffold formation is physically linked to the centrioles (8). Likewise, with the exception of the mitochondria(29), the segregation of most other organelles is also linked to the centrosome. Thus, we assessed whether the knockdown of TgCEP250L1 impacts on the segregation of the centrosome, and its associated structures. We observed that, while nuclear segregation is clearly aberrant upon TgCEP250L1 knockdown, the apicoplast is segregated to the daughter cells (**Fig. 1G**, asterisks indicate apicoplast DNA and **Fig. 2D** and inset). Likewise, the outer core of the centrosome, marked by the presence of TgCentrin1, is able to segregate into newly formed cells. Each daughter cell displays a single TgCentrin1 dot in physical proximity to the daughter cells’ apical ends, however TgCentrin1 is detectable distanced from the nucleus (**Fig. 2F**). Consistently, TEM analysis showed nuclei within the structure known as the residual body (30) (**Fig. 3A**). The residual body can be clearly distinguished from the daughter cells, as the former is only delimited by mother-cell derived plasma membrane, while the latter display the inner-membrane complex (**Fig. 3A**). Unpacked nuclei within the residual body are also observed by IFA, by labeling the plasma membrane with the marker TgSag1(**Fig. 3B**).

**Figure 3.**
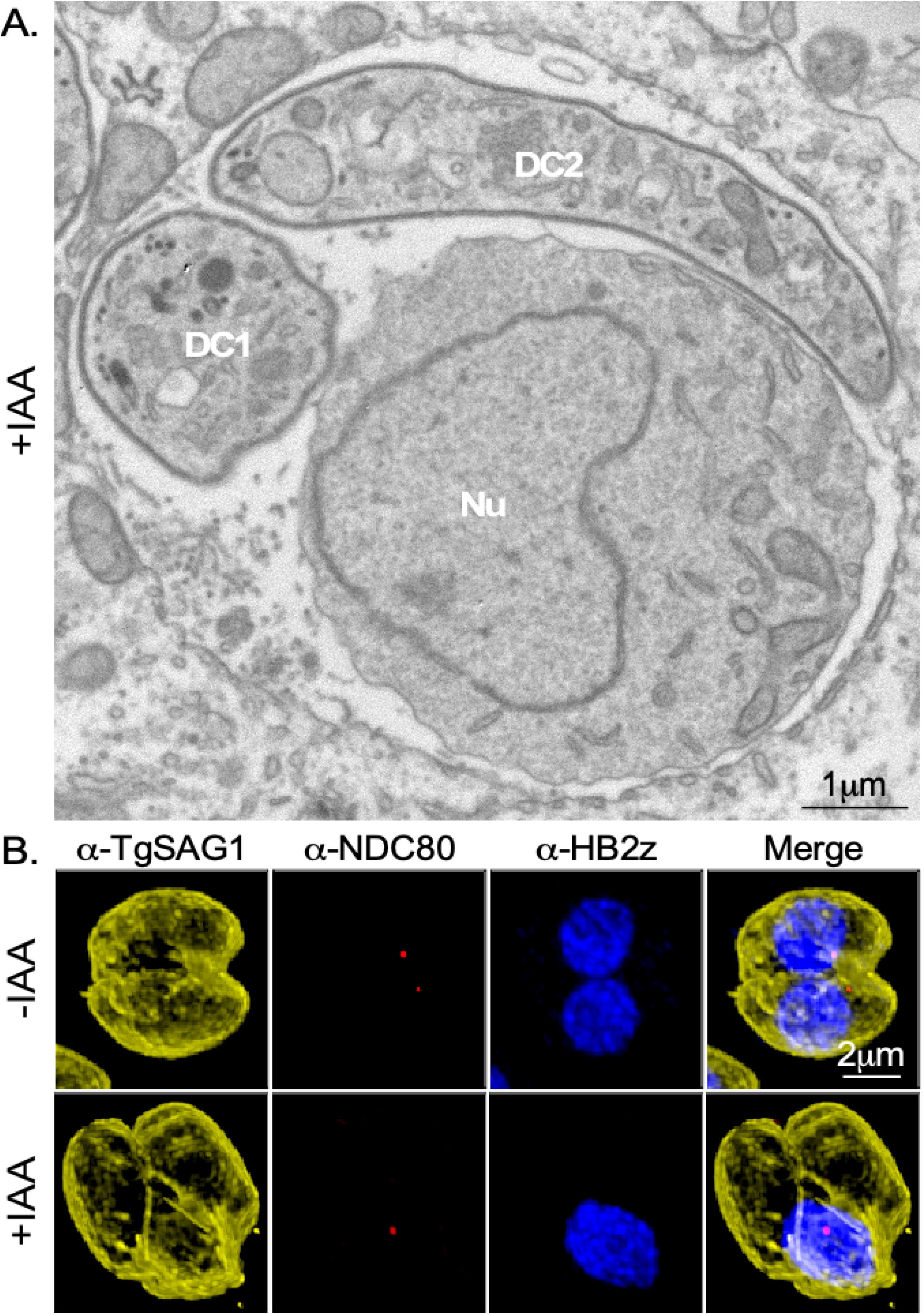
TgCEP250L1 knockdown causes nuclear loss to the residual body. **(A**) Electron micrograph of parasites treated with auxin for 24hs shows that **an unsegregated nucleus resides within the residual body after *T. gondii* division.** Note that two individual cells have been formed, but lack nuclei. Simultaneously, a large nucleus (Nu) stays in the residual body, recognized at the ultrastructural level by its lack of inner membrane complex. **(B**) Immunofluorescence assay of TgCEP250L1-mAID-HA parasites, stained with anti-NDC80, anti-H2Bz, and anti-SAG1, and treated as indicated. Note that the plasma membrane marker SAG1 labels the periphery of the residual body.

To gain insight into the molecular mechanism underlying the nuclear missegregation phenotype of the TgCEP250L1 knockdown cells, we set out to understand the dynamics of TgCEP250L1 with respect to structures known to impact nuclear segregation. The centrosome of *T. gondii* is at the limit of optical microscopy resolution (200 nm in size approx.). While much has been deciphered about the ultrastructure of the organelle using TEM, molecular insight into relative position of different elements (such as for example the inner and outer cores) has been gained through the use of super-resolution microscopy. A cost-effective alternative to the latter is the combined use of isotropic expansion and classical optical microscopy (confocal). Ultrastructure expansion microscopy (U-ExM) has been recently incorporated for use in *T. gondii* (31). In our hands, this technique allowed us expand parasites ~2.5-3.5 fold. We labeled expanded parasites with anti-acetylated tubulin, which labels the mitotic spindle, the centrioles and the cortical microtubules (**Fig. 4A**). The increase in resolution allowed us to clearly visualize that TgCEP250L1 does not co-localize with the centrioles. This is consistent with the premise that centrioles likely reside within the outer core (coinciding with the localization of TgCentrin1). We observe that TgCEP250L1 is dynamic throughout the cell cycle (**Fig. 4**). In recently divided parasites, early in interphase, TgCEP250L1 localizes adjacent to the centrioles (arrowheads, **Fig. 4**). We consistently observe that the TgCEP250L1 “dot” is asymmetrically distributed with respect to the centrioles’ position, being closer to one of the centrioles within the pair than to the other (**Fig. 4**, “cytokinesis/early interphase”). Upon entry into S-phase centrioles duplicate and the duplicated pairs start moving away from each other (28, 32). Shortly after, daughter cell scaffolds become apparent (7, 8). During the progression of mitosis, TgCEP250L1 evolves from a dot positioned in between the duplicated centriole pair, to an oval-like shape whose distance to the centrioles increases as daughter cells grow towards the mother cell periphery (**Fig. 4**). Eventually, the oval breaks into two distinct dots, each localizing at the tip of a tubulin-labeled structure, closely resembling the microtubule arrays of the mitotic spindle.

**Figure 4.**
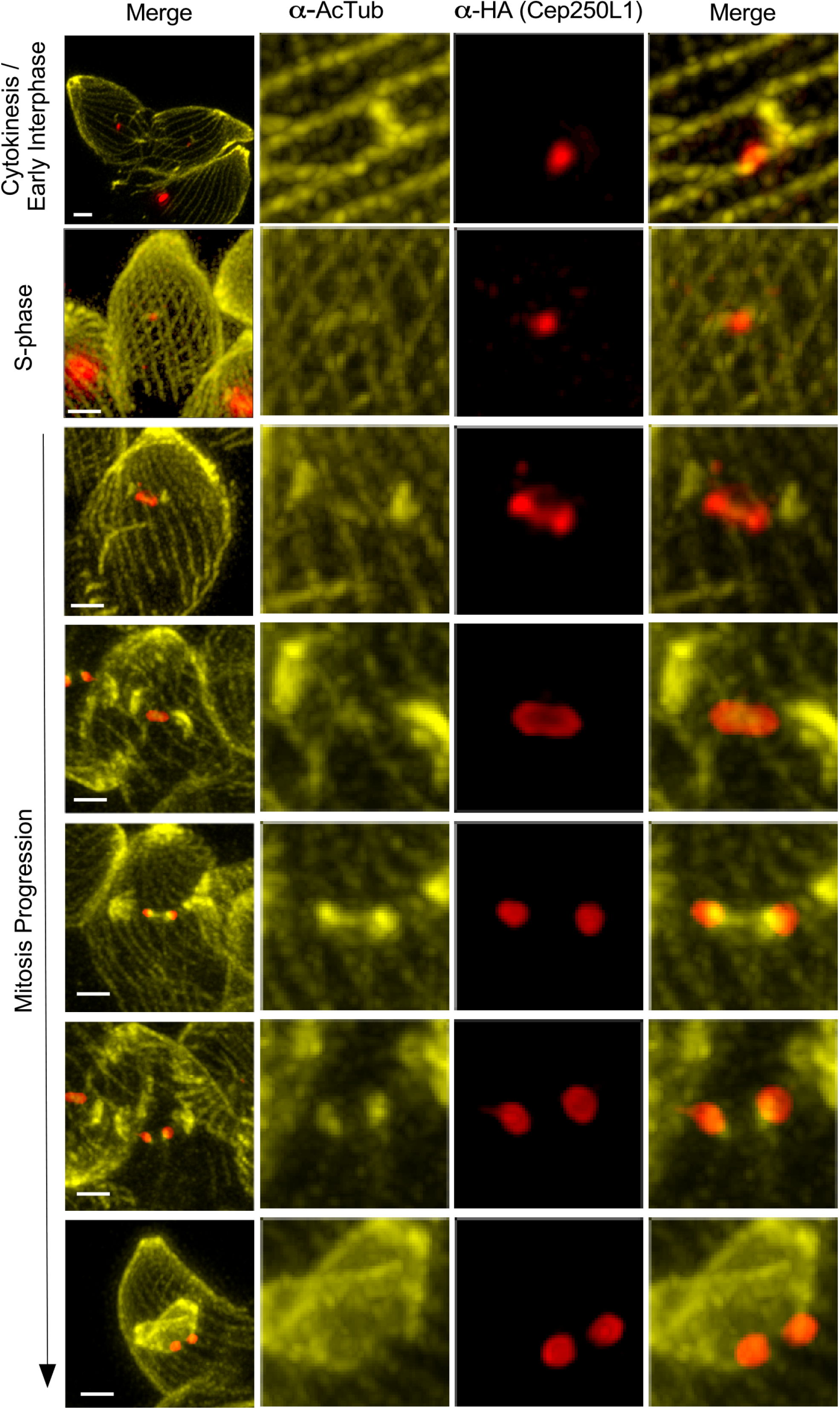
TgCEP250L1 displays a dynamic localization along the cell cycle linked to the position of the mitotic spindle. Ultrastructure expansion (U-ExM) rendered parasites 2.5-3.5 fold-larger. Parasites were labeled with antibodies as indicated. Note that acetylated tubulin is present in the scaffold microtubules of both the mother and the forming daughter cells, the centrioles of the centrosome, and the mitotic spindle. Scale bar represents 1mm in all cases. Arrowheads indicate the position of individual centrioles.

The mitotic spindle is a key “molecular machine” participating in chromosome segregation, made up of dynamic microtubules, microtubule binding proteins, and microtubule-bound motors. In *T. gondii*, spindle microtubules are only polymerized during cell division, and are absent during interphase (14). Very few molecular players involved in spindle assembly or stabilization have been identified in *T. gondii* (17, 33, 34). EB1s directly bind and stabilize microtubules in species ranging from yeast to humans; TgEB1 is a well conserved member of the EB1 protein family. It was previously shown that TgEB1 displays a dynamic localization along the cell cycle, residing in the nucleoplasm outside of division, concentrating adjacent to the centrosome in S-phase, and finally moving along with the spindle as cell division progresses (17). Using UExM, we visualize the relative positions of TgCEP250L1, TgEB1 and the mitotic spindle microtubules (**Fig. 5**). We observed that TgCEP250L1 localizes in between duplicated centriole pairs prior to the time when TgEB1 is detectable at the spindle. At the time when TgCEP250L1 adopts its “oval” staining pattern, TgEB1 is appreciably detectable both beneath the TgCEP250L1 oval and at the same site as TgCEP250L1. As TgCEP250L1 re-localizes to the proximal tip of the mitotic spindle microtubules asters, TgEB1 localizes beneath it, coinciding with the position of microtubules, as expected (**Fig. 5**, “mitosis progression”).

**Figure 5.**
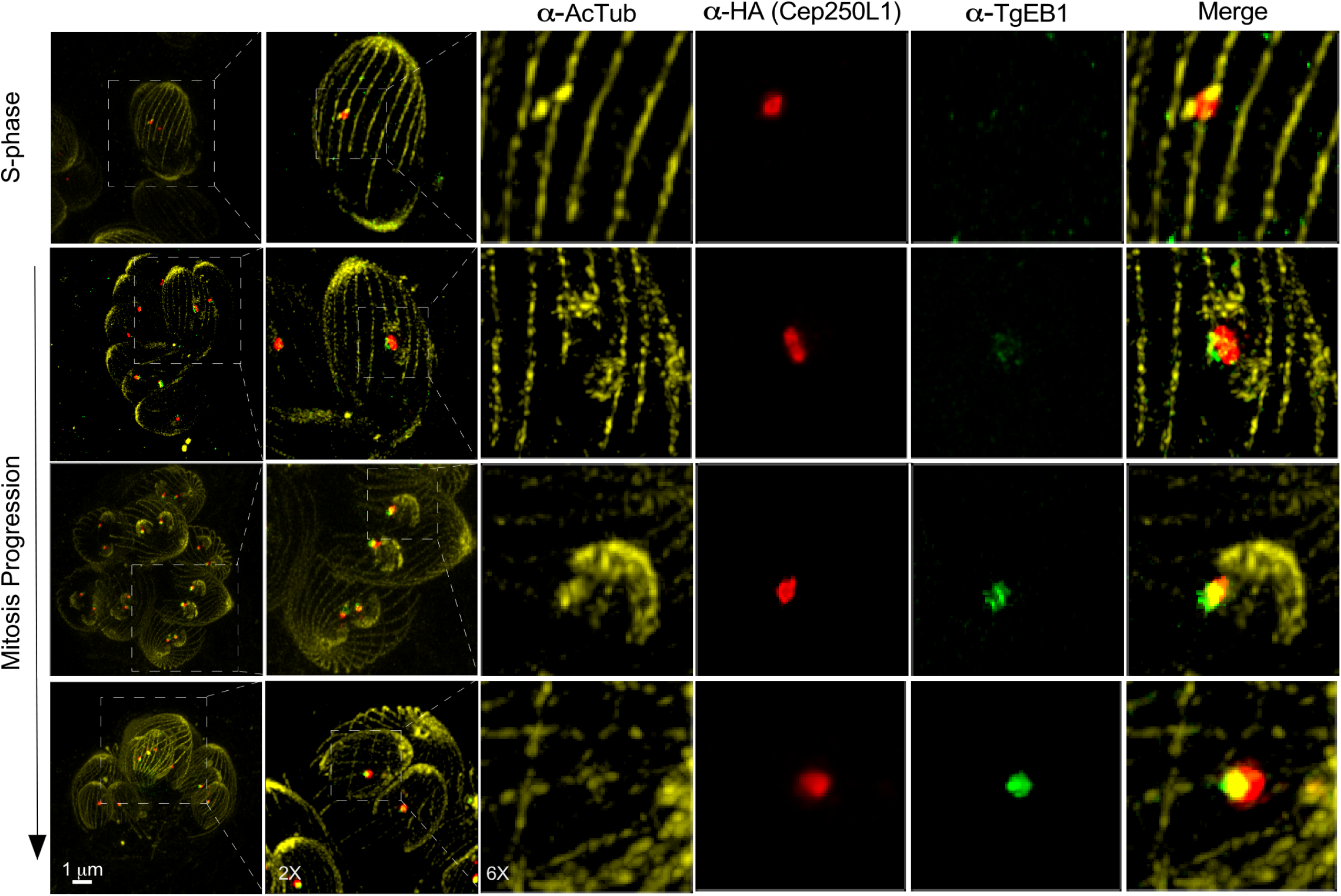
TgCEP250L1 dynamics correlates with that of a microtubule binding protein of the mitotic spindle. Ultrastructure expansion (U-ExM) rendered parasites 2.5-3.5 fold-larger. Parasites were labeled with antibodies as indicated. Note that acetylated tubulin is present in the scaffold microtubules of both the mother and the forming daughter cells, the centrioles of the centrosome, and the mitotic spindle. TgEB1 is a mitotic spindle marker which displays a dynamic cell-cycle dependent pattern of localization (17).

We then asked whether TgEB1’s localization to the mitotic spindle is affected upon TgCEP250L1’s knockdown. In non-dividing untreated parasites (-IAA), TgEB1 is virtually undetectable (**Fig. 6A**, -IAA upper panel). In stark contrast, dividing parasites (duplicated centrosomes labeled with anti-TgCentrin1, **Fig. 6A**, -IAA lower panel) TgEB1 markedly localizes in between the centrosomes, where the mitotic spindle resides. We observed that in parasites displaying the mutant phenotype (i.e. fragmented or unsegregated nuclei), TgEB1 is roughly detectable only in the nucleoplasm, with greater intensity in parts of the nucleus that appear to have fragmented (**Fig. 6A**, +IAA upper panel). However, we never detect it re-localizing to the expected mitotic spindle position, even in parasites bearing multiple TgCentrin1 dots (**Fig. 6A**, +IAA lower panel). The lack of TgEB1 recruitment to the location whereby the mitotic spindle is expected to assemble, correlates with lack of kinetochore segregation in dividing parasites (**Supplementary Fig. 3**).

**Figure 6.**
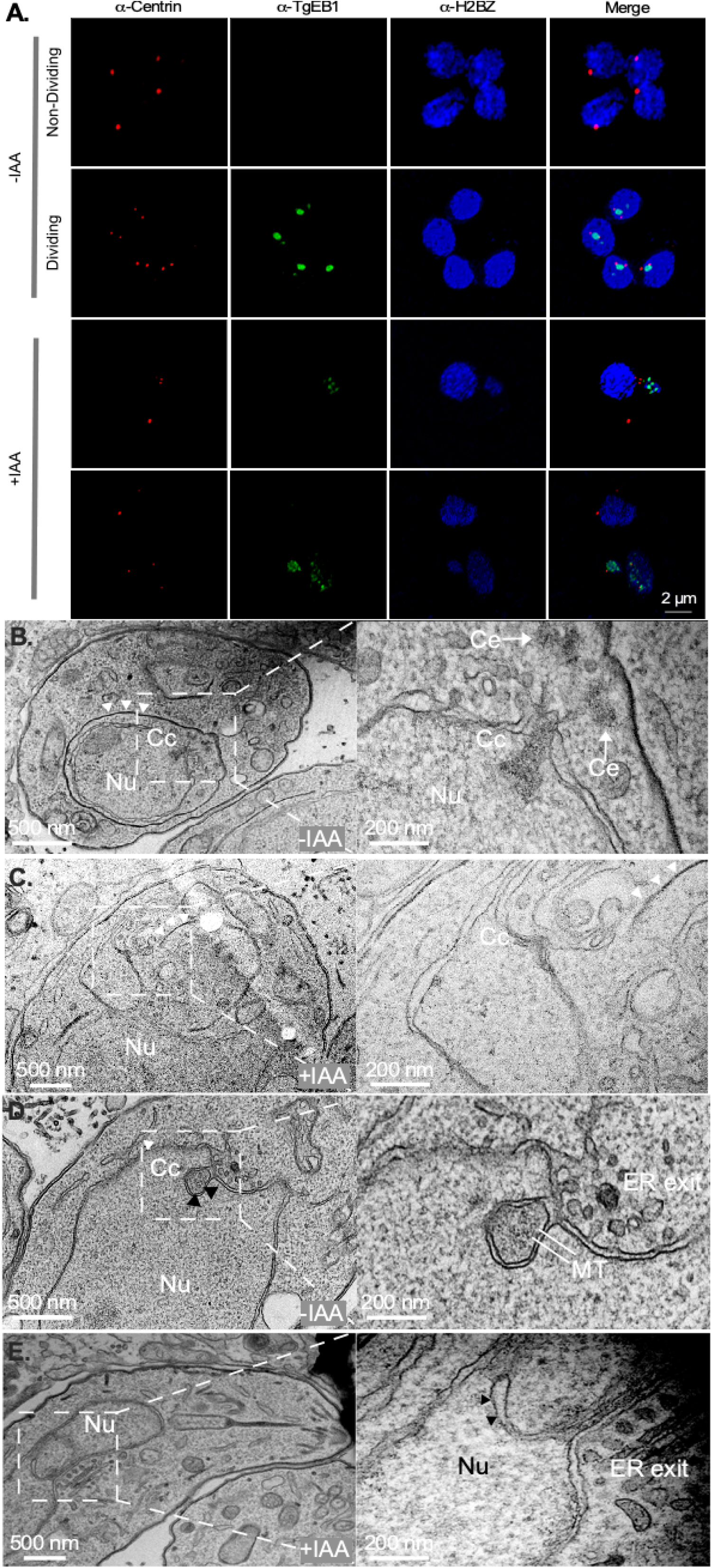
TgCEP250L1 knockdown causes mitotic spindle assembly defects. (A) Knockdown of TgCEP250L1 precludes TgEB1 translocation to the spindle. TgCEP250L1-mAID-HA parasites were stained with antibodies, and treated as indicated. In the presence of TgCEP250L1 untreated parasites exhibit either largely undetectable TgEB1 outside of division (upper panel) or readily detectable TgEB1 in between the duplicated centrosomes (red) during division (lower panel). The latter corresponds with the expected position of the mitotic spindle. In treated parasites, whereby TgCEP250L1 is knocked down, TgEB1 remains in the nucleoplasm when detectable (both panels) and does not re-localize even in the presence of multiple centrosomes, indicative of cell division (lower panel). (**B) The centrocone houses the mitotic spindle during cell division**. Transmission Electron micrograph of untreated TgCEP250L1-mAID-3HA parasites (transversal cut) shows the centrioles (Ce), centrocone (Cc) and segregating nucleus (Nu). Note that the centrocone position in the nuclear envelope is always adjacent to the ER exit site. White arrowheads indicate the inner membrane complex position of the assembling daughter cell. (**C) TgCEP250L1 knockdown alters the centrocone structure**. An electron micrograph (longitudinal view) of an IAA treated parasite reveals a bilobed large nucleus, physically distant from the site of daughter cell assembly (white arrowheads indicate the inner membrane complex). Note that adjacent to the ER exit site, an elaboration of the nuclear envelope, which is possibly a reminiscent of the centrocone structure, can be appreciated. (**D) The mitotic spindle is housed within the nuclear envelope in untreated parasites**. Electron micrograph of untreated TgCEP250L1-mAID-3HA parasites shows that the centrocone (Cc) houses the mitotic spindle microtubules (MT) within the nuclear envelope (black arrowheads) and is adjacent to the ER exit site. **(E) TgCEP250L1 knockdown collapses the centrocone structure**. Electron micrograph of parasites treated with IAA for 24hs. Note that a void invagination of the nuclear envelope can be seen adjacent to the ER exit site. Mitotic spindle microtubules are undetectable.

The mitotic spindle is known to assemble adjacent to the centrosome, within a conical elaboration of the nuclear envelope, devoid of chromatin, known as the centrocone (35–37). Figure 6B shows a nucleus at the end of karyokinesis/mitosis showing a centrocone with mitotic spindle adjacent to centrioles. Conspicuously, by TEM, we observed that parasites which fail to segregate their nuclei display protrusions of the nuclear envelope in the vicinity of the ER exit site (**Fig. 6C**), the region of the nuclear envelope whereby the centrocone (Cc) and the spindle normally form (**Fig. 6A and D**). However, this protrusions only vaguely resembles the centrocone morphology during division, as it is devoid of detectable spindle microtubules (inset of **Fig. 6C**). Mitotic spindle microtubules can be detected in normally dividing parasites within invaginations of the nuclear envelope (**Fig. 6D**, arrows in inset). We detect nuclear envelope invaginations in IAA treated adjacent to the ER exit site (**Fig. 6E**). However, these are devoid of detectable microtubules.

γ-tubulin is a crucial protein participating in spindle microtubule nucleation through the nucleation of the γ-tubulin ring complex (γTuRC). However, γ-tubulin has been shown to localize at the outer core of the centrosome (9), a location that, *a priori*, is physically distant from the site of mitotic spindle nucleation and of the previously reported TgCEP250L1 location at the inner core. We reckon that if the inner core is involved in the mitotic spindle microtubules dynamics, γ-tubulin should be detected at this site. We evaluated the localization of γ-tubulin with respect to TgCEP250L1 and observed that, in line with previous reports, γ-tubulin co-localizes with the centrioles at the centrosomal outer core (9). However, we also detected it at a distant location from the centrioles, co-localizing with the mitotic spindle and TgCEP250L1, during mitosis (**Fig. 7A**).

**Figure 7.**
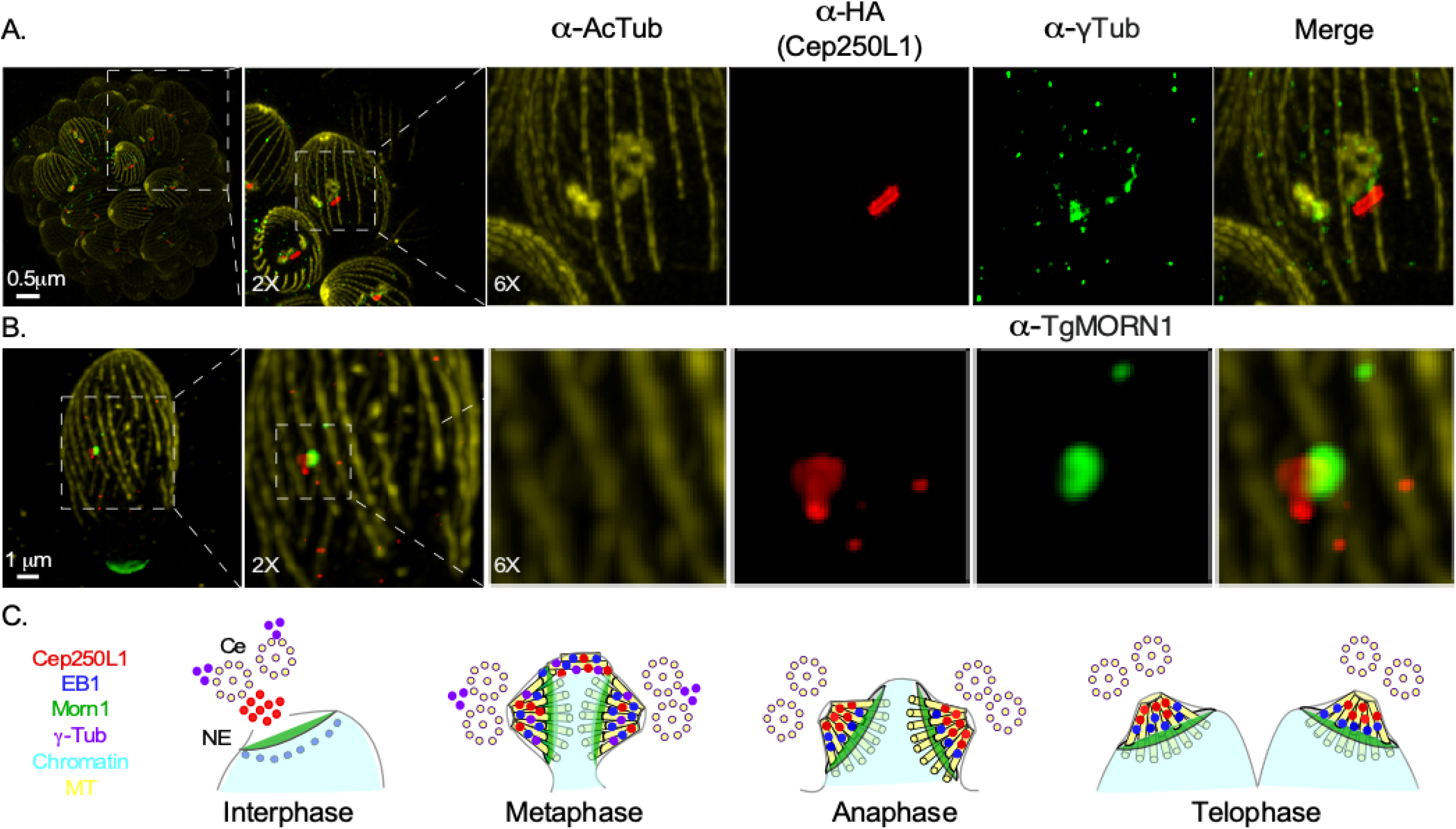
Ultrastructure expansion microscopy (U-ExM) shows that TgCEP250L1 co-localizes with γ-tubulin and the centrocone marker TgMORN1. (**A**) Dividing parasites were labeled with antibodies as indicated. Note that during division TgCEP250L1 partially colocalizes with γ-tubulin and the mitotic spindle. (**B) TgCEP250L1 partially co-localizes with the centrocone marker TgMORN1**. Note that TgCEP250L1 (red) partially colocalizes with TgMORN1 (green). Parasites were labeled as indicated. (**C) Schematic representation of the TgCEP250L1 localization dynamics along the cell cycle progression**. A schematic representation of the relative localization of TgCEP250L1 along the cell cycle, with respect to the indicated structures, is shown.

The centrocone is marked at its base by TgMORN1(36), we reckon that if the inner core is involved in the formation or stability of the mitotic spindle, then its localization should be intimately related to that of the centrocone. Using UExM, we observed that TgCEP250L1 is detectable physically adjacent and partially co-localizing with TgMORN1 both in interphase and in dividing parasites (**Fig. 7B**). To gauge whether TgCEP250L1 could be within the centrocone, we compiled our TEM micrographs and determined that the centrocone spans approximately 250-350 nm from its base (where TgMORN1 resides) to its tip. We established in our UExM experiments that TgCEP250L1 is detectable from right on top of TgMORN1 to 300 nm outwards, further reinforcing the notion that TgCEP250L1 could reside within the nuclear elaboration of the centrocone, where the mitotic spindle nucleates during mitosis (**Fig. 7C**).

## DISCUSSION

Animal centrosomes display a protein called c-NAP1 which participates in physically linking centrioles. Phosphorylation of c-NAP1 catalyzes the dissolution of the linkage between mother and daughter centrioles. This process is catalyzed prior to mitosis by a NIMA-related kinase, called Nek1, to allow the two new centrosomes, duplicated during S-phase, to eventually migrate to opposite poles of the cell. A conditional mutant of TgNek1 results in parasites displaying a single centrosome, suggesting that this kinase regulates centrosome splitting in an akin manner to what happens in animals (7, 10). *T. gondii’s* genome was initially proposed to encode two homologs of c-NAP1, which were identified principally based on the presence of coiled-coiled domains; TgCEP250 and TgCEP250L1 (9). However, *in silico* studies proposed that neither of the two proteins are *bona fide* homologs of cNAP-1 (38). In accordance, it has been experimentally proven that TgCEP250 is not a substrate of TgNek1(10). On the other hand, TgCEP250L1 locates at the inner core, distant from the outer core where centrioles reside (9).

In this work we described the function and dynamic localization of TgCep250L1, the sole marker of the inner core as a proxy of the inner core function. We observed that upon depletion of TgCep250L1, parasites exhibit nuclear segregation defects, while normally forming daughter cells. Moreover, the centrosome - as defined by centriolar markers - disconnects from the nucleus, suggesting that TgCep250L1 plays an essential role in coordinating nuclear and centrosome co-segregation into forming daughter cells. We further describe the dynamic localization of TgCep250L1 as the cell cycle, and particularly, as spindle formation progresses. We uncovered that TgCep250L1 displays a highly dynamic localization pattern, which particularly correlates with the dynamics of the microtubule binding protein EB1 and mitotic spindle formation.

It was established in the early 60s, by transmission electron microscopy, that a mitotic spindle assembles adjacent to the centrosome in *T. gondii* (4, 39). Microtubules remain extranuclear, residing within a conical elaboration of the nuclear envelope; a structure coined more recently as “the centrocone” (40). Previous studies have addressed the dynamics of the spindle, showing that recruitment of tubulin to the centrocone happens at the end of interphase/G1, just before the onset of S-phase, a stage marked by the duplication of the centrosome (17, 28, 32, 36). The chromosomes, through their centromeres and their associated kinetochore proteins, are associated with the centrocone throughout the cell cycle, but are only reached by the growing spindle during mitosis (14, 20, 25). Although the ultrastructure of the mitotic spindle, and its dynamics, have been broadly described, the molecular pathways involved in the nucleation of the microtubule in *T. gondii* remain poorly understood.

The PCM serves as a platform for protein complexes that regulate organelle trafficking, protein degradation and spindle assembly in animal cells. This complex and dynamic matrix of proteins visibly envelope the centrioles. A vast number of protein components of the PCM have been identified in species ranging from humans to flies. These include, but are not limited to: CEP152, CEP57, CPAP (SAS-4), γ-tubulin, CEP192, pericentrin, CDK5RAP2 (Cnn) and a number of microtubule binding/stabilizing proteins, together with regulatory kinases and phosphatases (including Polo-like kinase 1, Aurora A kinase, and PP2 homologs). Strikingly, many of the functionally most important PCM proteins seem to be absent from the *T. gondii* genome. Proteins like CEP152, CEP192, PLK1-4, pericentrin and Cnn are not coded for in this parasite (9, 38, 41).

*T. gondii* γ-tubulin had been previously localized to the outer core of the centrosome, a location where centrioles reside (9, 42). However, spindle microtubule nucleation does not occur at this location but rather at the opposite core of the centrosome. Here, we show that during mitosis, γ-tubulin displays a dual localization both to the outer core but also, as expected, to the inner core. This is consistent with its expected role in MT nucleation of the mitotic spindle, but also consistent with the fact that MTs of the cortical cytoskeleton of the daughter cell are nucleated at the outer core.

A number of kinases, including a NIMA-related Kinase, Cyclin dependent-related kinases (CRK), Aurora-related kinases, and mitogen-activated protein-related kinases (MAPKs) have been shown to impact centrosome biology in *T. gondii*. TgMAPKL1 temperature sensitive mutants display over duplication of Centrin1 with respect to TgCEP250L1, suggesting that this protein plays a role in negatively regulating outer core biogenesis - and indirectly demonstrating that the inner and outer cores are separately regulated(9). On the contrary, a conditional knockdown of MAPK2 displays under-duplicated Centrin1 and TgCEP250L1, suggesting that the presence of this kinase positively regulates the duplication of both cores of the centrosome(43). Centrosome position was found altered upon knock down of TgCDPK7 (44). Work by Alvarez and collaborators showed that depletion of Cycling-related kinase CRK6 impairs the duplication of the centrocone and generates defects in kinetochores segregation (45). Three serine/threonine Aurora related kinases are functionally related to licensing events of the cell cycle(33, 46). Depletion of TgArk-1 leads to defects in the duplication of both the spindle pole and the inner core (33). TgArk-2 localizes at the intranuclear mitotic spindle. However, mutants of TgArk-2 are able to proliferate normally (46). TgArk-3 localizes at the outer core of the centrosome and has been linked with regulation of the budding process (46).

More recently, a homolog of MAD1, a component of the spindle assembly checkpoint (SAC), was shown to localize at the *T. gondii* kinetochore, suggesting a canonical pathway of mitotic fidelity checkpoint could operate and be conserved in apicomplexans (47). The canonical notion of checkpoint, however, is in contrast with the phenotype observed in null mutants of the kinetochore protein NDC80, whereby the budding part of the cell cycle progresses in spite of the fact that kinetochores cannot attach to the spindle. Consistent with our proposition that TgCEP250L1 is involved in mitotic spindle assembly or stability, the knockdown of Ndc80 renders parasites with a very similar phenotype to that of the TgCEP250L1 mutant shown here, whereby nuclei fail to segregate but daughter cells assemble nonetheless (20).

The modular nature of cell division in *T. gondii* (i.e. nuclear division and budding operate through seemingly independent regulatory networks) allows this parasite to divide using distinct modes, including endodyogeny and schizogony. This is also at the basis of pathogenicity, as it allows this parasite to modulate the output of its replication and adapt to specific cell niches. Endodyogeny and schizogony have traditionally been described as differing only in whether cytokinesis follows nuclear mitosis immediately or not. In endodyogeny, nuclear mitosis is immediately followed by budding. In contrast, in schizogony, budding only occurs following multiple rounds of nuclear mitosis. And the dual organization of the *T. gondii* centrosome has been proposed to contribute to this cell division flexibility, whereby the outer core determines the number of daughter cells assembled while the inner core controls nuclear events, separately.

Endodyogeny has often been referred to as “small scale” schizogony. Nonetheless, our deep understanding of schizogonic cell division comes from studying it in the *Plasmodium* genus. However, central structural differences exist between *T. gondii* and *Plasmodium* that raise questions about the conservation of certain regulatory principles between the schizogony in these species. One such difference is that *T. gondii* displays microtubule-based centrioles, while *Plasmodium* species do not. *Plasmodium falciparum* nuclear MTOCs have been referred to as the centriolar plaque (CP). Until very recently, it was considered that the CP resided within the nuclear envelope, much like the budding yeast spindle pole body does. However, new advances in microscopy (including the combination of UExM with STED and CLEM) allowed Simon and collaborators to observe and describe a bipartite structure whereby an extranuclear segment houses markers such as Centrin, and an intranuclear segment (albeit devoid of chromatin) houses tubulin and presumably other unidentified proteins involved in microtubule nucleation (48). The centrioles themselves in *T. gondii* have been shown to critically regulate the formation of daughter cells through the assembly of a fiber that positions the apical MTOC(8). However, how microtubules of the mitotic spindle are assembled in either species has only recently begun to be addressed. Work by Simon and collaborators helped clarify that microtubule nucleation during schizogonic mitosis occurs at an intranuclear site. Concomitantly, Liffner and Absalon showed that the spindle undergoes dynamic changes which accompany the chromatin dynamics as mitosis progresses (49). Non-dividing nuclei sustain a handful (~5) of bundled individual intranuclear microtubules which have been named the “hemi-spindle.” This spindle extends from the single CP to the opposite side of the nucleus. Upon DNA replication onset in S-phase, the hemi-spindle retracts, the CP duplicates, and the mitotic spindle forms. The spindle connects to the chromosome’s kinetochores (47), while keeping the CPs interconnected by spanning the nucleus forming the “interpolar spindle” (49). The differences in mitotic spindle dynamics correlated with differences in kinetochore connections with the spindle. Kinetochores are constitutively attached to the centrocone in *T. gondii*, while there seem to be lateral connections to the chromosomes, in addition to connections to the kinetochores, in *Plasmodium*.

Overall, these new molecular and structural insights suggest that while the nature of the nuclear MTOCs in *T. gondii* and *Plasmodium* species is different (i.e. centrioles vs. no centrioles), the organization principle is indeed highly similar. For the organization of their nuclear mitosis, both species rely on a PCM-like protein cumulate - of yet to be deciphered, albeit likely divergent, composition - physically distant from *bona fide* centrosomal markers such as centrin, which reside within specialized structures - centrioles or the CP distal to the nuclear envelope. Nonetheless, subtle differences exist; the *Plasmodium’s* PCM-like proteins are intranuclear while *T. gondii*’s TgCEP250L1, EB1 and the microtubules stay distal to the edge of the nuclear envelope marked by TgMORN1, suggesting these structures are mostly extranuclear.

Our results support the function of the inner core as a key element for organization or stability of the mitotic spindle, and further support the notion that a dual-organized centrosome is the basis of compartmentalizing functions providing cell cycle flexibility in *T. gondii*. Future studies addressing the cell-cycle specific interactions of TgCEP250L1 and their interplay to regulate the mitotic spindle will shed light on the detailed mechanisms of action of this protein and of the specialized region of the nuclear periphery it inhabits.

## ACKNOWLEDGEMENTS

The authors gratefully acknowledge the Cell Biology Unit, UMPI; the joint research unit between the Institut Pasteur de Montevideo and the Instituto Nacional de Investigación Agropecuaria (INIA), the Advanced Bioimaging Unit at the Institut Pasteur Montevideo for their support & assistance in the present work. We would like to especially thank M.Sc. Marcela Diaz, Tabare de los Campos and M. Sc. Paula Céspedes. We thank the TEM Unit at the School of Sciences, Universidad de la República. We are especially grateful to Dr. Gabriela Casanova, Dr. Gaby Martínez and M.Sc. Magela Rodao for their diligent assistance in acquisition and processing of images. We also thank the TEM units of the Center of Microscopy at the Universidade Federal de Minas Gerais, Belo Horizonte, and the Centro Nacional de Biologia Estrutural e Bioimagem, Rio de Janeiro, Brazil. We thank Dr. Gonzalo Ferreira and Ms. Mariana Di Doménico for assistance with image processing. RT and MEF are PEDECIBA researchers. MEF is an SNI researcher.

## REFERENCES

1. Adl SM, Simpson AGB, Lane CE, Lukeš J, Bass D, Bowser SS, Brown MW, Burki F, Dunthorn M, Hampl V, Heiss A, Hoppenrath M, Lara E, Gall L Le, Lynn DH, McManus H, Mitchell EAD, Mozley-Stanridge SE, Parfrey LW, Pawlowski J, Rueckert S, Shadwick L, Schoch CL, Smirnov A, Spiegel FW. 2012. The revised classification of eukaryotes. J Eukaryot Microbiol 59:429–514.

2. De Lima Bessa G, Wagner De Almeida Vitor R, Erica ·, Martins-Duarte S. Toxoplasma gondii in South America: a differentiated pattern of spread, population structure and clinical manifestations. Parasitol Res 1:3.

3. Commodaro AG, Belfort RN, Rizzo LV, Muccioli C, Silveira C, Burnier MN, Belfort R. 2009. Ocular toxoplasmosis - An update and review of the literature. Mem Inst Oswaldo Cruz https://doi.org/10.1590/S0074-02762009000200030.

4. van den Zypen E, Piekarski G. 1968. Ultraestructura de la endodiogenia en Toxoplasma gondii. Bol Chil Parasitol 23:90–94.

5. Morrissette N, Gubbels M-J. 2020. The Toxoplasma cytoskeleton: structures, proteins, and processesToxoplasma gondii. LTD.

6. Francia ME, Striepen B. 2014. Cell division in apicomplexan parasites. Nat Rev Microbiol.

7. Chen CT, Gubbels MJ. 2013. The Toxoplasma gondii centrosome is the platform for internal daughter budding as revealed by a Nek1 kinase mutant. J Cell Sci https://doi.org/10.1242/jcs.123364.

8. Francia ME, Jordan CN, Patel JD, Sheiner L, Demerly JL, Fellows JD, de Leon JC, Morrissette NS, Dubremetz J-F, Striepen B. 2012. Cell Division in Apicomplexan Parasites Is Organized by a Homolog of the Striated Rootlet Fiber of Algal Flagella. PLoS Biol 10:e1001444.

9. Suvorova ES, Francia M, Striepen B, White MW. 2015. A Novel Bipartite Centrosome Coordinates the Apicomplexan Cell Cycle. PLoS Biol 13.

10. Chen CT, Gubbels M. 2019. TgCep250 is dynamically processed through the division cycle and is essential for structural integrity of the Toxoplasma centrosome. Mol Biol Cell https://doi.org/10.1091/mbc.E18-10-0608.

11. Courjol F, Gissot M. 2018. A coiled-coil protein is required for coordination of karyokinesis and cytokinesis in Toxoplasma gondii. Cell Microbiol 20:e12832.

12. Brown KM, Long S, Sibley LD. 2017. Plasma Membrane Association by N-Acylation Governs PKG Function in Toxoplasma gondii. MBio 8.

13. Brown K, Long S, Sibley L. 2018. Conditional Knockdown of Proteins Using Auxin-inducible Degron (AID) Fusions in Toxoplasma gondii. BIO-PROTOCOL https://doi.org/10.21769/bioprotoc.2728.

14. Francia ME, Bhavsar S, Ting LM, Croken MM, Kim K, Dubremetz JF, Striepen B. 2020. A Homolog of Structural Maintenance of Chromosome 1 Is a Persistent Centromeric Protein Which Associates With Nuclear Pore Components in Toxoplasma gondii. Front Cell Infect Microbiol https://doi.org/10.3389/fcimb.2020.00295.

15. Sidik SM, Hackett CG, Tran F, Westwood NJ, Lourido S. 2014. Efficient Genome Engineering of Toxoplasma gondii Using CRISPR/Cas9. PLoS One 9:e100450.

16. Mann T, Gaskins E, Beckers C. 2002. Proteolytic processing of TgIMC1 during maturation of the membrane skeleton of Toxoplasma gondii. J Biol Chem 277:41240–41246.

17. Chen CT, Kelly M, De Leon J, Nwagbara B, Ebbert P, Ferguson DJP, Lowery LA, Morrissette N, Gubbels MJ. 2015. Compartmentalized Toxoplasma EB1 bundles spindle microtubules to secure accurate chromosome segregation. Mol Biol Cell 26:4562–4576.

18. Bogado SS, Dalmasso MC, Ganuza A, Kim K, Sullivan Jr WJ, Angel SO, Vanagas L. 2014. Canonical histone H2Ba and H2A.X dimerize in an opposite genomic localization to H2A.Z/H2B.Z dimers in Toxoplasma gondii. Mol Biochem Parasitol 197:36–42.

19. Vanagas L, Jeffers V, Bogado SS, Dalmasso MC, Sullivan Jr WJ, Angel SO. 2012. Toxoplasma histone acetylation remodelers as novel drug targets. Expert Rev Anti Infect Ther 10:1189–1201.

20. Farrell M, Gubbels MJ. 2014. The Toxoplasma gondii kinetochore is required for centrosome association with the centrocone (spindle pole). Cell Microbiol 16:78–94.

21. Dos Santos Pacheco N, Soldati-Favre D. 2021. Coupling Auxin-Inducible Degron System with Ultrastructure Expansion Microscopy to Accelerate the Discovery of Gene Function in Toxoplasma gondii, p. 121–137. In de Pablos, LM, Sotillo, J (eds.), Parasite Genomics: Methods and Protocols. Springer US, New York, NY.

22. Martins-Duarte ES, Portes J de A, da Silva RB, Pires HS, Garden SJ, de Souza W. 2021. In vitro activity of N-phenyl-1,10-phenanthroline-2-amines against tachyzoites and bradyzoites of Toxoplasma gondii. Bioorg Med Chem 50:116467.

23. Hu K, Roos DS, Murray JM. 2002. A novel polymer of tubulin forms the conoid of Toxoplasma gondii. J Cell Biol https://doi.org/10.1083/jcb.200112086.

24. Köhler S, Delwiche CF, Denny PW, Tilney LG, Webster P, Wilson RJM, Palmer JD, Roos DS. 1997. A Plastid of Probable Green Algal Origin in Apicomplexan Parasites. Science (80-) 275:1485–1489.

25. Brooks CF, Francia ME, Gissot M, Croken MM, Kim K, Striepen B. 2011. Toxoplasma gondii sequesters centromeres to a specific nuclear region throughout the cell cycle. Proc Natl Acad Sci U S A 108:3767–3772.

26. Behnke MS, Wootton JC, Lehmann MM, Radke JB, Lucas O, Nawas J, Sibley LD, White MW. 2010. Coordinated progression through two subtranscriptomes underlies the tachyzoite cycle of toxoplasma gondii. PLoS One https://doi.org/10.1371/journal.pone.0012354.

27. Striepen B, Crawford MJ, Shaw MK, Tilney LG, Seeber F, Roos DS. 2000. The plastid of Toxoplasma gondii is divided by association with the centrosomes. J Cell Biol https://doi.org/10.1083/jcb.151.7.1423.

28. Verhoef JMJ, Meissner M, Kooij TWA. 2021. Organelle dynamics in apicomplexan parasites. MBio 12.

29. Ovciarikova J, Lemgruber L, Stilger KL, Sullivan WJ, Sheiner L. 2017. Mitochondrial behaviour throughout the lytic cycle of Toxoplasma gondii. Sci Rep https://doi.org/10.1038/srep42746.

30. Attias M, Miranda K, De Souza W. 2019. Development and fate of the residual body of Toxoplasma gondii. Exp Parasitol 196:1–11.

31. Tosetti N, Pacheco N dos S, Bertiaux E, Maco B, Bournonville L, Hamel V, Guichard P, Soldati-Favre D. 2020. Essential function of the alveolin network in the subpellicular microtubules and conoid assembly in Toxoplasma gondii. Elife https://doi.org/10.7554/eLife.56635.

32. Hartmann J, Hu K, He CY, Pelletier L, Roos DS, Warren G. 2006. Golgi and centrosome cycles in Toxoplasma gondii. Mol Biochem Parasitol 145:125–127.

33. Berry L, Chen CT, Francia ME, Guerin A, Graindorge A, Saliou JM, Grandmougin M, Wein S, Bechara C, Morlon-Guyot J, Bordat Y, Gubbels MJ, Lebrun M, Dubremetz JF, Daher W. 2018. Toxoplasma gondii chromosomal passenger complex is essential for the organization of a functional mitotic spindle: a prerequisite for productive endodyogeny. Cell Mol Life Sci 75:4417–4443.

34. Naumov A, Kratzer S, Ting L-M, Kim K, Suvorova ES, White MW. 2017. The Toxoplasma Centrocone Houses Cell Cycle Regulatory Factors. MBio https://doi.org/10.1128/mbio.00579-17.

35. Dubremetz JF. 1973. Etude ultrastructurale de la mitose schizogonique chez la coccidie Eimeria necatrix (Johnson 1930). J Ultrastruct Res 42:354–376.

36. Gubbels MJ, Vaishnava S, Boot N, Dubremetz JF, Striepen B. 2006. A MORN-repeat protein is a dynamic component of the Toxoplasma gondii cell division apparatus. J Cell Sci 119:2236–2245.

37. Dubremetz J-F. 1971.. L’ultrastructure du centriole et du centrocone chez la coccidie eimeria necatrix. É tude au cours de la schizogonie 23:453–458.

38. Morlon-guyot J, Francia ME, Dubremetz J, Wassim D. 2017. Towards a molecular architecture of the centrosome in Toxoplasma gondii 55–71.

39. Sheffield HG, Melton ML. 1968. The fine structure and reproduction of Toxoplasma gondii. J Parasitol https://doi.org/10.2307/3276925.

40. Dubremetz J. 1971. L’ultrastructure du centriole et du centrocone chez la coccidie eimeria necatrix. Étude au cours de la schizogonie. J Microsc 23:453–8.

41. Tomasina R, González FC, Francia ME. 2021. Structural and Functional Insights into the Microtubule Organizing Centers of Toxoplasma gondii and Plasmodium spp. Microorganisms 9:2503.

42. Morrissette N. 2015. Targeting Toxoplasma tubules: Tubulin, microtubules, and associated proteins in a human pathogen. Eukaryot Cell https://doi.org/10.1128/EC.00225-14.

43. Hu X, O’shaughnessy WJ, Beraki TG, Reese ML. 2020. Loss of the conserved alveolate kinase MAPK2 decouples toxoplasma cell growth from cell division. MBio https://doi.org/10.1128/mBio.02517-20.

44. Bansal P, Antil N, Kumar M, Yamaryo-Botté Y, Rawat RS, Pinto S, Datta KK, Katris NJ, Botté CY, Keshava Prasad TS, Sharma P. 2021. Protein kinase TgCDPK7 regulates vesicular trafficking and phospholipid synthesis in Toxoplasma gondii. PLOS Pathog 17:e1009325.

45. Alvarez CA, Suvorova ES. 2017. Checkpoints of apicomplexan cell division identified in Toxoplasma gondii. PLoS Pathog https://doi.org/10.1371/journal.ppat.1006483.

46. Berry L, Chen CT, Reininger L, Carvalho TG, El Hajj H, Morlon-Guyot J, Bordat Y, Lebrun M, Gubbels MJ, Doerig C, Daher W. 2016. The conserved apicomplexan Aurora kinase TgArk3 is involved in endodyogeny, duplication rate and parasite virulence. Cell Microbiol https://doi.org/10.1111/cmi.12571.

47. Brusini L, Santos Pacheco N Dos, Soldati-Favre D, Brochet M. 2021. Organization and composition of apicomplexan kinetochores reveal plasticity in chromosome segregation across parasite modes of division. bioRxiv 2021.11.03.466924.

48. Simon CS, Voβ Y, Funaya C, Machado M, Penning A, Klaschka D, Cyrklaff M, Kim J, Ganter M, Guizetti J. 2021. An extended DNA-free intranuclear compartment organizes centrosomal microtubules in Plasmodium falciparum. bioRxiv https://doi.org/10.1101/2021.03.12.435157.

49. Liffner B, Absalon S. 2021. Expansion Microscopy Reveals Plasmodium falciparum Blood-Stage Parasites Undergo Anaphase with A Chromatin Bridge in the Absence of Mini-Chromosome Maintenance Complex Binding Protein. Microorganisms 9.

